# Artificial mutations in the nuclear gene encoding mitochondrial RNA polymerase restore pollen fertility in cytoplasmic male sterile tomato

**DOI:** 10.1101/2025.02.18.638787

**Authors:** Kosuke Kuwabara, Rika Nakajima, Alexis Gaetan Van Bosstraeten, Kentaro Ezura, Kinya Toriyama, Tohru Ariizumi, Kenta Shirasawa

**Affiliations:** Graduate School of Agricultural Science, Tohoku University, Sendai, Miyagi, 980-8572, Japan; Graduate School of Life and Environmental Sciences, University of Tsukuba, Tsukuba, Ibaraki 305-8572, Japan; Graduate School of Science and Technology Degree Programs in Life and Earth Sciences Master’s Program in Agro-Bioresources Science and Technology, University of Tsukuba, Ibaraki 305-8577, Japan; Institute of Life and Environmental Sciences, University of Tsukuba, Tsukuba, Ibaraki 305-8572, Japan; Tsukuba Plant Innovation Research Center, Tsukuba, Ibaraki 305-8577, Japan; Kazusa DNA Research Institute, Kisarazu, Chiba 292-0818, Japan

**Keywords:** *Solanum lycopersicum*, Restorer of fertility, Cytoplasmic male sterility, Mutagenesis, Hybrid seed production, Mitochondrial RNA polymerase

## Abstract

Cytoplasmic male sterility (CMS) and restorer of fertility (*Rf*) are important traits in F_1_ hybrid breeding. However, the CMS/*Rf* system has not been used in tomatoes because of the limited resources of *Rf* lines. In this study, we performed mutagenesis in tomato CMS seeds and successfully obtained 13 suppressor mutants with pollen fertility. Using bulked segregant analysis and whole-genome sequencing for each suppressor mutant, we detected mutations associated with fertility restoration in the nuclear-encoded gene for the mitochondrial RNA polymerase termed *SlRPOTm* in four independent mutants created through mutagenesis. Furthermore, we found that the loss of function of *SlRPOTm* was associated with fertility restoration in the tomato CMS line. Expression analysis of *orf137*, a tomato CMS-causing gene, revealed that reduced expression of *orf137* was associated with fertility restoration in tomato CMS. In addition, F_1_ plants carrying mutations in *SlRPOTm* were generated, and tomato fruit formation was comparable to that of normal F_1_ plants. This study demonstrates for the first time that the loss of function of mitochondrial RNA polymerase contributes to fertility restoration in CMS lines. Furthermore, it is possible to replace various tomato varieties with *Rf* lines using genome editing technology, which will promote tomato F_1_ breeding in the future.

## Introduction

Most commercial vegetables and field crops are F_1_ hybrids because they produce a significant increase in crop yield and provide stability in food production (Duvick, 1999). F_1_ hybrid breeding has long been used for tomatoes (*Solanum lycopersicum*), one of the most widely cultivated and consumed vegetables worldwide. However, the considerable amount of labor and the maintenance of high seed purity have been major issues in hybrid seed production (Priyadarsini *et al*., 2025).

Cytoplasmic male sterility (CMS) is caused by incompatibility of mitochondrial and nuclear genes, resulting in abnormal male organ function. It has been detected in more than 150 species of higher plants (Bohra *et al*., 2016). CMS plants do not produce seeds through self-pollination, thus CMS is valuable for hybrid breeding because it eliminates the need to remove anthers and maintains high hybrid seed purity. Efficient hybrid seed production using CMS has been implemented in several species, including rice, sunflowers, sorghum, maize, and radish (Bohra *et al*., 2016). CMS lines have been artificially generated in tomato using recurrent backcrossing and transgenic experiments (Petrova *et al*., 1999; Sandhu *et al*., 2007). Furthermore, asymmetric cell fusion between cultivated tomato lines and the wild potato relative *S. acaule* produced CMS tomato plants (Melchers *et al*., 1992). Recently, we identified *orf137* in the mitochondrial genome as a CMS-causing gene in these tomato CMS lines (Kuwabara *et al*., 2021, 2022). However, in vegetables that are fertilized to produce fruit, such as tomatoes, both CMS lines and *restorer of fertility* (*Rf*) genes encoded in the nuclear genome are required to restore pollen fertility (Chen and Liu, 2014). Three wild relatives, *S. pimpinellifolium* (LA1670), *S. lycopersicum* var. *cerasiforme* (LA1673), and *S. cheesmaniae* (LA0166) have *Rf* genes that restore fertility in CMS lines possessing *orf137* (Watabe *et al*., 1997). We recently identified multiple gene loci for fertility restoration in these wild relatives; however, fertility restoration by these relatives is weak (Iki *et al*., 2025). Therefore, novel *Rf* lines are required to apply the CMS/*Rf* system to tomato breeding.

Several wild plant species have *Rf* genes in their nuclear genomes, which repress the expression of CMS-causing genes in the mitochondrial genome. In many cases, PPR proteins induce fertility restoration through RNA processing or the degradation of CMS-causing gene products (Chen and Liu, 2014). However, it was recently reported that mutagenic breeding produces *Rf* lines in *Arabidopsis thaliana*. Seeds of the *Arabidopsis* CMS line possessing *orf117Sha* as a CMS-causing gene were treated with ethyl methanesulfonate (EMS), a reagent that induces random mutations, to obtain several suppressors (or called fertile revertants). A loss-of-function mechanism of the PPR protein in these suppressor mutants has been previously discussed (Durand *et al*., 2021). Furthermore, the *Rf* gene identified in Chinese wild rice (CW)-type CMS encodes *RETROGRADE-REGULATED MALE STERILITY* and a single nucleotide polymorphism (SNP) in the promoter determines fertility restoration (Fujii and Toriyama, 2009). These studies suggested that EMS, which causes random base substitutions, is a useful tool for generating novel *Rf* lines in various plant species.

In the present study, we aimed to develop tomato *Rf* lines using EMS mutagenesis. Seeds of tomato CMS lines were treated with EMS, and 13 suppressor mutants were successfully developed. Bulked segregant analysis (BSA) identified mutations in the mitochondrial RNA polymerase termed *SlRPOTm*, and genome editing technology demonstrated that the loss of function of *SlRPOTm* restores pollen fertility in tomato CMS lines. Furthermore, mutation of *SlRPOTm* induced sufficient fruit production in tomato F_1_ lines. This study provides the first evidence that mitochondrial RNA polymerase is involved in fertility restoration and contributes to the promotion of F_1_ hybrid breeding using the CMS/*Rf* system in tomatoes.

## Results

### Development of new tomato Rf lines by EMS mutagenesis

EMS treatments of seeds of the tomato CMS line Dwarf “CMS[P]” were conducted twice. The first treatment was 0.5% (v/v) EMS for Dwarf “CMS[P]” seeds, and approximately 600 plants of the M_1_ generation were grown. In the second treatment, 1% (v/v) EMS was used on Dwarf “CMS[P]” seeds, and approximately 1,200 plants of the M_1_ generation were grown. suppressor mutants were screened based on whether the plants produced seeds via self-pollination. A total of 13 suppressor mutants were obtained: three suppressor mutants in the first treatment (termed #1, #2, and #7) and ten suppressor mutants in the second treatment (termed #9, #10, #11, #12, #14, #16, #17, #20, #21, and #22). To maintain each suppressor mutant and to exclude mutations that were not associated with fertility restoration, suppressor mutants were continuously backcrossed to Dwarf “CMS[P]” (Figure 1). Pollen germination phenotypes were examined in the BC_2_F_1_ generation. As these lines were backcrossed to wild-type (WT) Dwarf “CMS[P],” the mutations related to fertility restoration were expected to be heterozygous. Germinated pollen rates varied among the lines, with the highest rate of 39% in EMS#1 and the lowest rate of 2% in EMS#2. Because tomato CMS pollen exhibits expansion or burst at the apertures (i.e., germination pores) (Kuwabara *et al*., 2022), we also counted these abnormal pollens. Most suppressor mutants showed decreased rates of abnormal pollen, but the rates of EMS#17 and #22 were the same as that of Dwarf “CMS[P]” (Figure 2a, b). In summary, pollen germination was observed in 13 suppressor mutants, and the proportion of abnormal pollen decreased in many lines, confirming reliable pollen fertility in these lines.

**Figure 1.**
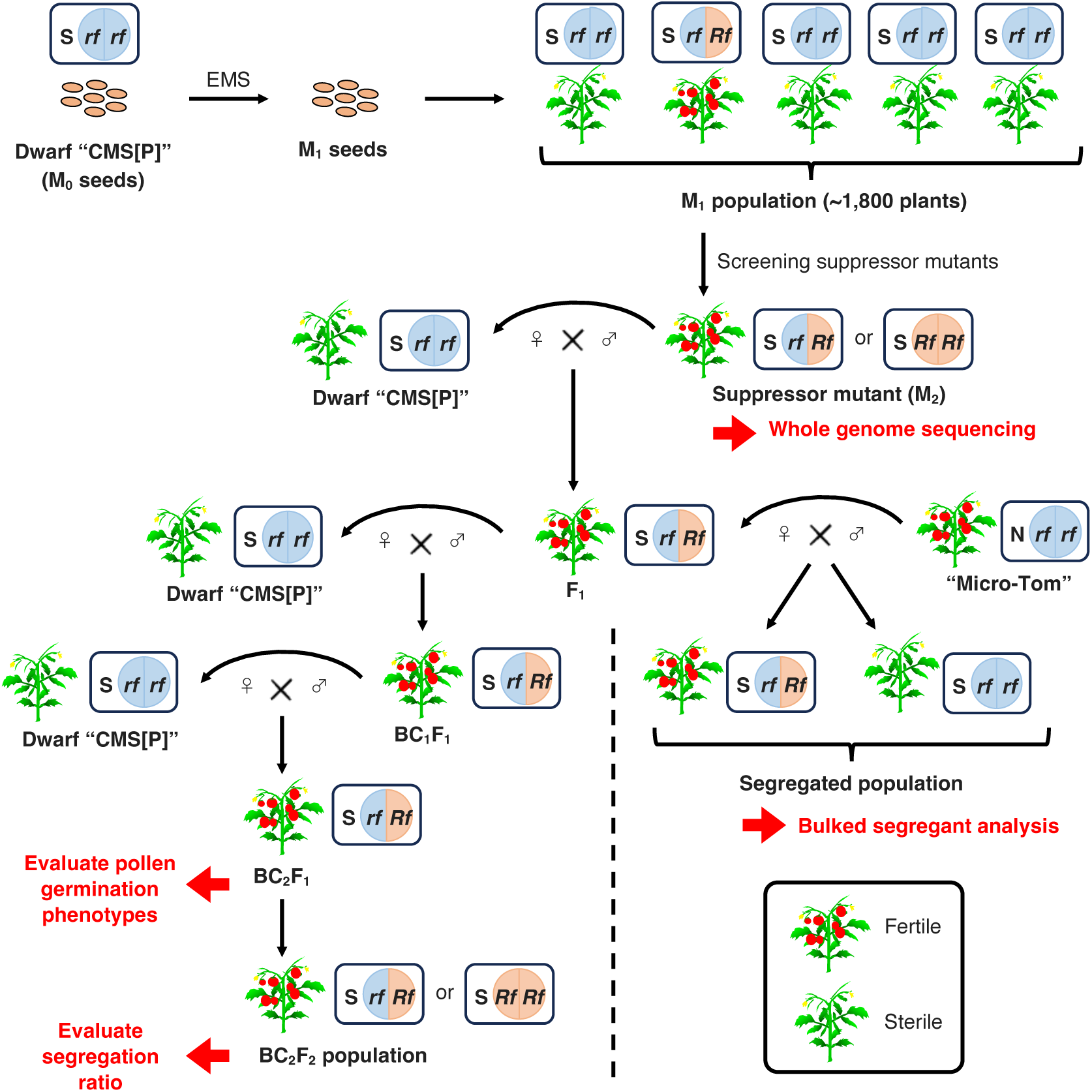
EMS mutagenesis to develop suppressor mutants in tomato CMS line. Seeds of Dwarf “CMS[P]” (M_0_ seeds) were treated with EMS to yield M_1_ (EMS-treated) seeds. M_1_ plants were grown and seed formation was evaluated by self-pollination. Only the M_1_ plants with seed formation were selected as suppressor mutants. To exclude mutations that are not associated with fertility restoration, mutants were continuously backcrossed to Dwarf “CMS[P]” to create BC_2_F_1_ generation and used for evaluation of pollen germination phenotypes. These plants were also self-pollinated to create BC_2_F_2_ populations to evaluate segregation ratios. For bulked segregant analysis, F_1_ plant of EMS#1 was cross-pollinated with pollen of “Micro-Tom” and a segregated population with fertile and sterile plants was produced. S: sterile cytoplasm; Rf: restorer gene (or mutation)-associated fertility restoration; rf: nonfunctional restorer gene.

**Figure 2.**
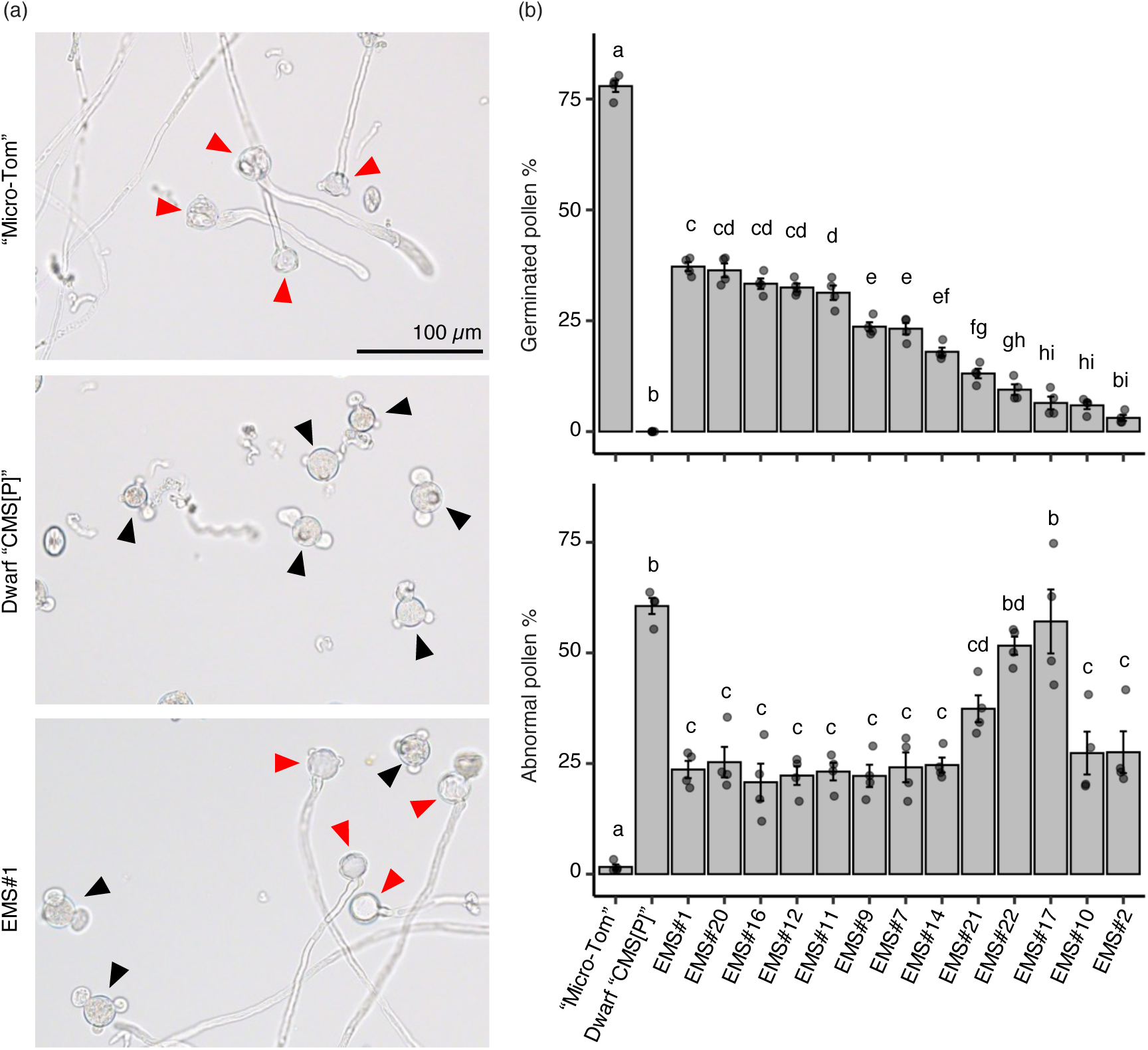
Pollen phenotypes in suppressor mutants. (a) Phenotypes of pollen after 4 h of incubation in liquid germination media for “Micro-Tom”, Dwarf “CMS[P]”, and BC_2_F_1_ plant of EMS#1. Red triangles indicate germinated pollen, and black triangles indicate abnormal (triple-tips or burst) pollen. (b) The percentage of germinated pollen (top panel) and abnormal pollen (bottom panel). Data were obtained from four independent experiments (n = 4). Different letters above the bars indicate statistically significant differences, as determined by Tukey’s HSD test (P < 0.05). Error bars represent standard errors.

### Mutations of Solyc05g010660 are candidates for fertility restoration

In this study, EMS#1 was subjected to BSA to identify the mutations responsible for fertility restoration. We cross-pollinated WT “Micro-Tom” pollen into the F_1_ generation of EMS#1 to obtain a segregated population (Figure 1). In this population, 13 individuals were pollen-fertile, and 21 were sterile. We performed whole-genome sequencing on bulk DNA of fertile and sterile populations, detected SNPs, and calculated the ΔSNP-index. Although no significant peaks were detected in the ΔSNP-index plot, a weak peak was detected around 5 Mb on chromosome 5 (Figure S1); therefore we focused the region between 2 to 8 Mb on this chromosome. We set two criteria to narrow down the SNPs related to fertility restoration in EMS#1: (1) 0.3 ≤ SNP-index ≤ 0.7 in fertile bulk and SNP-index = 0 in sterile bulk; (2) SNPs in intergenic regions and synonymous SNPs were excluded. Based on these criteria, two SNPs on *Solyc05g009410* and *Solyc05g010660* were identified. Furthermore, we conducted whole-genome sequencing of the M_2_ generation of each suppressor mutant and detected mutations of *Solyc05g010660* in EMS#7, #9, and #11, while no mutations of *Solyc05g009410* were detected in all mutants. In *Solyc05g010660*, a mutation was detected in intron 6 in EMS#1, a nonsense mutation appeared at amino acid 657 in EMS#7, a splicing mutation occurred at the splice donor site of intron 11 in EMS#9, and a missense mutation (p.Gly608Ser) occurred in EMS#11 (Figure 3a). To investigate the segregation pattern of these mutations, BC_2_F_1_ plants with heterozygous mutations were self-pollinated, and BC_2_F_2_ populations were generated (Figure 1). Owing to the gametophytic mode of fertility restoration in the tomato CMS line (Iki *et al*., 2025), all BC_2_F_2_ individuals were expected to possess a causal mutation. As expected, all BC_2_F_2_ individuals from EMS#1, #7, #9, and #11 harbored mutations (Table 1), suggesting that these mutations in *Solyc05g010660* are candidates for fertility restoration.

**Figure 3.**
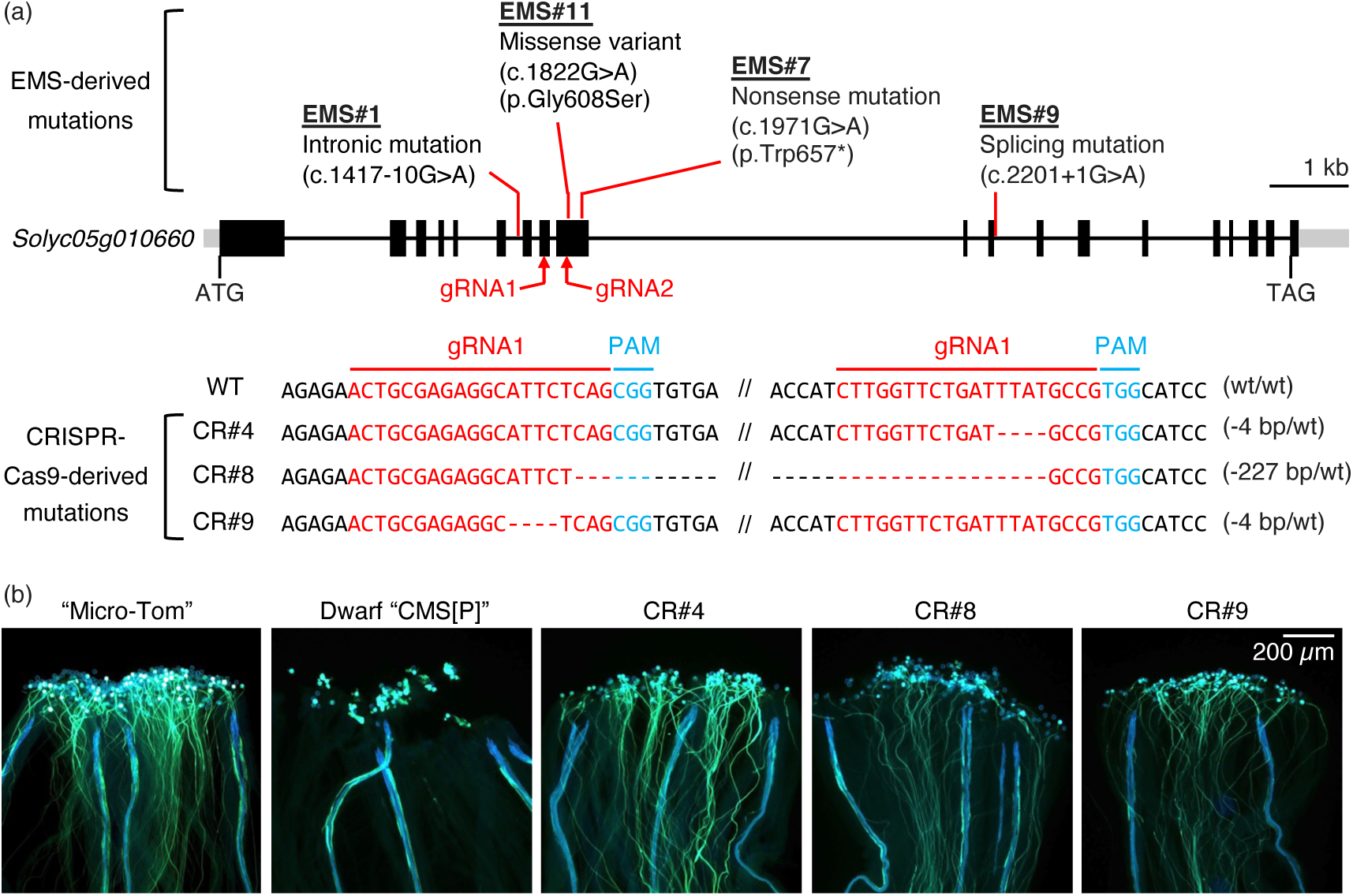
Artificial mutations of the *Solyc05g010660* gene restore pollen fertility in tomato CMS line. (a) Schematic representation of the *Solyc05g010660* gene structure and induced mutations. Mutations derived from EMS mutagenesis and CRISPR-Cas9 are shown. The positions of guide RNAs (gRNA1 and gRNA2) used for CRISPR-Cas9 editing are indicated in red, and the protospacer adjacent motif (PAM) sequences are shown in blue. CRISPR-Cas9-induced mutations were detected in T_0_ plants. The symbol “-” denotes deleted nucleotides. wt: wild-type. (b) Pollen germination phenotypes were observed 24 hours after pollination in Micro-Tom, Dwarf “CMS[P]”, and CRISPR-Cas9-generated mutants (CR#4, CR#8, CR#9) of *Solyc05g010660*. Pollen tubes were stained with aniline blue. The fluorescent lines observed in Dwarf “CMS[P]” represent vascular bundles.

**Table 1.**
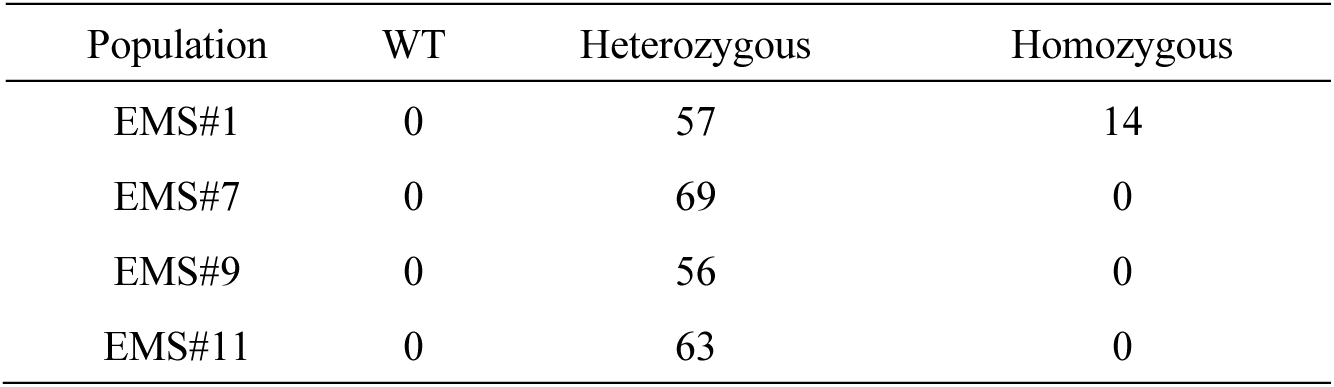
Genetic analysis of *rpoTm* mutations in BC_2_F_2_ population.

*Solyc05g010660* has high homology with the nuclear-encoded T7 phage-type RNA polymerase that localizes to the mitochondria in *Arabidopsis* (*AtRPOTm*). To confirm whether the protein encoded by *Solyc05g010660* localized to the mitochondria, it was fused with superfolder Green Fluorescent Protein (sfGFP) at the C-terminus and transiently expressed in the protoplasts of *Nicotiana benthamiana*. GFP fluorescence of the fused protein was consistent with the fluorescence of the mitochondrial marker MitoTracker Red but did not overlap with the autofluorescence of chloroplasts (Figure S2). This result demonstrated that Solyc05g010660 which is homologous to AtRPOTm translocated into the mitochondria; therefore, we referred to it as *Solyc05g010660 SlRPOTm*.

### Loss of function of SlRPOTm restores pollen fertility in a tomato CMS line

EMS#7 and #9 have a nonsense mutation and a splicing mutation, respectively, suggesting that the loss of function of *Solyc05g010660* is related to fertility restoration. Therefore, we knocked out *SlRPOTm* in Dwarf “CMS[P]” using the CRISPR-Cas9 system and obtained three independent knockout lines (CR#4, CR#8, CR#9). Four-bp deletions were detected at different positions in CR#4 and CR#9, and a 227-bp deletion appeared in CR#8 as heterozygous mutations (Figure 3a). Both deletions caused frameshifts, suggesting that *SlRPOTm* was functionally defective. Pollen germination was observed on the stigma of three knockout lines, which was the same result in the fertile control line “Micro-Tom.” Pollen germination was not observed in the Dwarf “CMS[P]” (Figure 3b). These results demonstrate that the loss of function of *SlRPOTm* in tomato CMS lines contributes to fertility restoration.

### Effects of rpoTm mutation on male and female organs and plant growth

In *Arabidopsis*, the loss of function of *AtRPOTm* negatively affects female gametogenesis and embryogenesis; therefore, homozygous mutants cannot be obtained (Kühn *et al*., 2009; Tan *et al*., 2010). In the present study, homozygous individuals were not obtained from the BC_2_F_2_ populations of EMS#7, #9, and #11 (Table 1), suggesting that *rpoTm* mutations impaired female gametogenesis and embryogenesis in tomatoes, similar to the *Arabidopsis rpoTm* mutant. By contrast, homozygous individuals were obtained in the BC_2_F_2_ population of EMS#1, albeit at a lower ratio (Table 1). This result was demonstrated by both the CAPS marker and Sanger sequencing analyses (Figure S3a, b). EMS#1 has an intronic mutation 10 bp upstream of exon 7. To investigate how this mutation affected the splicing of *SlRPOTm* gene, we performed RT-PCR using primers targeting exons 6 and 7 and then performed Sanger sequencing. In the homozygous mutant of EMS#1, overlapping peaks were detected only between exons 6 and 7 (Figure S4a) and were divided by CRISP-ID (Dehairs *et al*., 2016). In addition to the cDNA from eight bases upstream of exon 7 the cDNA of WT *RPOTm* was also detected. This result suggests that G to A intronic mutation of EMS#1 forms a new splicing site “AG” and splicing occurs at both the new and original sites (Figure S4b). In other words, the intronic mutation in EMS#1 is expected to be a partial knockdown rather than a complete breakdown of *SlRPOTm* gene function. This may explain why homozygous individuals were obtained from EMS#1.

We further evaluated the effects of *rpoTm* mutants on plant growth using homozygous EMS#1. The homozygous mutant showed slower plant growth than WT “Micro-Tom,” Dwarf “CMS[P],” and heterozygous EMS#1 mutants (Figure S5a, b). Fresh weight was significantly lower in the homozygous mutants (Figure S5c). In addition, the number of days to flowering was delayed by 2 to 3 days in the homozygous mutants (Figure S5d). These results indicated that the homozygous mutation exerted negative effects on plant growth, whereas the heterozygous mutation had no effect.

Anthers and pollen were investigated to determine the effect of the EMS#1 mutation on male organs. There was no difference in flower appearance and anther width between “Micro-Tom,” Dwarf “CMS[P],” heterozygous and homozygous EMS#1 mutants. The pollen number was also unchanged in the homozygous mutants (Figure 4a, b). The pollen germination rate of EMS#1 homozygous mutants was 78%, which was similar to that of the fertile control (“Micro-Tom”). The pollen germination rate of the heterozygous mutant was 37%, whereas that of the homozygous mutant was only twice as high. The abnormal pollen rate was significantly reduced in homozygous mutants compared with those of heterozygous mutants and Dwarf “CMS[P]” (Figure 4c). These results indicate that the *rpoTm* mutation positively regulates pollen germination but does not negatively affect male organs.

**Figure 4.**
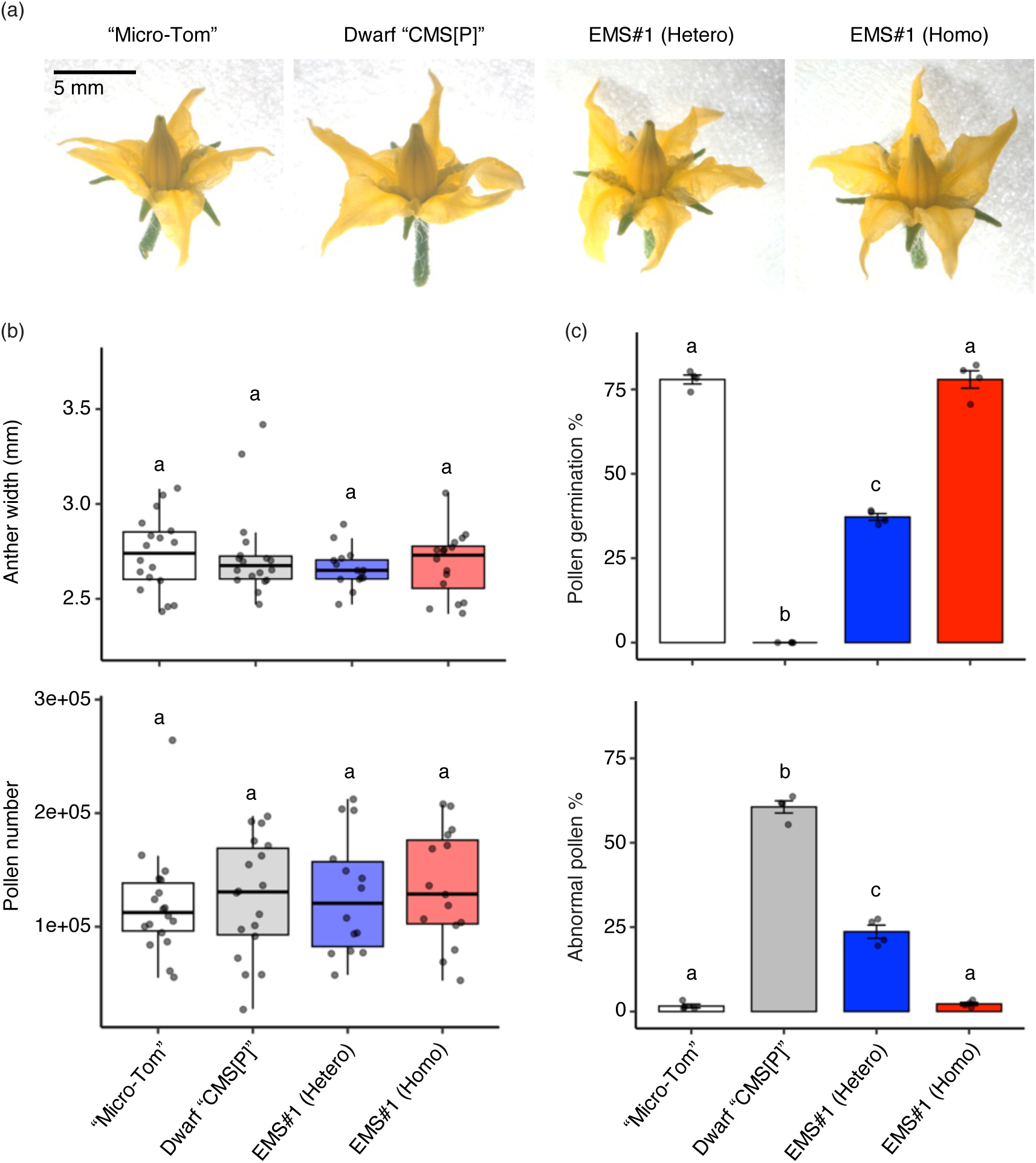
Phenotypes of male organs in heterozygous and homozygous mutants of EMS#1. (a) Flowers of “Micro-Tom”, Dwarf “CMS[P]”, EMS#1 heterozygous, and EMS#1 homozygous mutants. (b) Box plots showing anther width (top panel, n ≥ 15) and pollen number per flower (bottom panel, n ≥ 14) at the anthesis stage. The central line represents the median, the box edges indicate the interquartile range (IQR), and the whiskers extend to the minimum and maximum values within 1.5× IQR. Individual data points are shown as dots. Different letters indicate statistically significant differences (P < 0.05, Tukey’s HSD test). (c) Bar graphs showing the percentage of germinated pollen (top panel) and abnormal pollen (bottom panel) in liquid germination media (n = 4). Bars represent mean values, and error bars indicate standard error. Different letters indicate statistically significant differences (P < 0.05, Tukey’s HSD test).

### Association of lower expression level of orf137 with fertility restoration

In *Arabidopsis*, AtRPOTm has been shown to specifically recognize mitochondrial promoter sequences in vitro and accurately initiate transcription. Most mitochondrial promoters contain a YRTA motif (Y = T or C and R = A or G) near the transcription initiation sites (Hess and Börner, 1999; Kühn *et al*., 2007, 2005). To investigate whether this motif is present in the promoter region of *orf137*, which is a tomato CMS gene, the ends of *orf137* mRNA were determined using circularized reverse-transcription PCR (CR-RT-PCR). The search for the longest untranslated region (UTR) revealed that the 5’ UTR was 353 bp and the 3’ UTR was 106 bp. CGTA, one of the YRTA motifs, was detected 4 bp upstream of the 5’ UTR, suggesting that SlRPOTm recognizes this sequence and initiates transcription of *orf137* (Figure S6).

We next investigated the expression level of *orf137* using RT-qPCR. RNA was extracted from anthers at the anthesis stage, pollen incubated for 10 min (just before pollen germination), and pollen incubated for 60 min (after germination). The relative expression level of *orf137* in anthers and incubated pollen was significantly lower in homozygous EMS#1 compared to that in Dwarf “CMS[P]” (Figure 5a). We further performed RNA-Seq analysis in the anther and incubated pollen of homozygous mutant and Dwarf “CMS[P]” to evaluate the changes in the mitochondrial transcriptome. RNA-Seq reads were mapped to the mitochondrial genome of “CMS[P]” and calculated the mapping rates. The mapping rate to the mitochondria in pollen incubated for 10 and 60 min was approximately 1/50 of that in the anther (Table S1). These results were similar to the RT-qPCR results, which clearly showed a lower expression of *orf137* in pollen than in anthers, suggesting that more sophisticated techniques are needed to detect the mitochondrial transcriptome in pollen. Therefore, in this study, we focused on the RNA-Seq results of anthers. The mapping rate in homozygous EMS#1 was significantly lower than that in Dwarf “CMS[P]” anthers, suggesting that the total mitochondrial RNA content was reduced by the *rpoTm* mutation (Figure 5b, Table S1). The relative expression levels of most mitochondrial genes tended to decrease in the EMS#1 group, with significant reductions in *orf137*, *rps4*, *nad9*, *rrn26*, *nad6*, *nad5*, *rrn18,* and *atp1* (Figure 5c). Both RNA-Seq and RT-qPCR showed reduced mRNA levels of *orf137*. Given that *orf137* has been identified as a CMS-causing gene in the tomato CMS line (Kuwabara *et al*., 2022), reduced expression of this gene is likely linked to fertility restoration; however, decreased expression of other mitochondrial genes may be related to retardation of plant growth.

**Figure 5.**
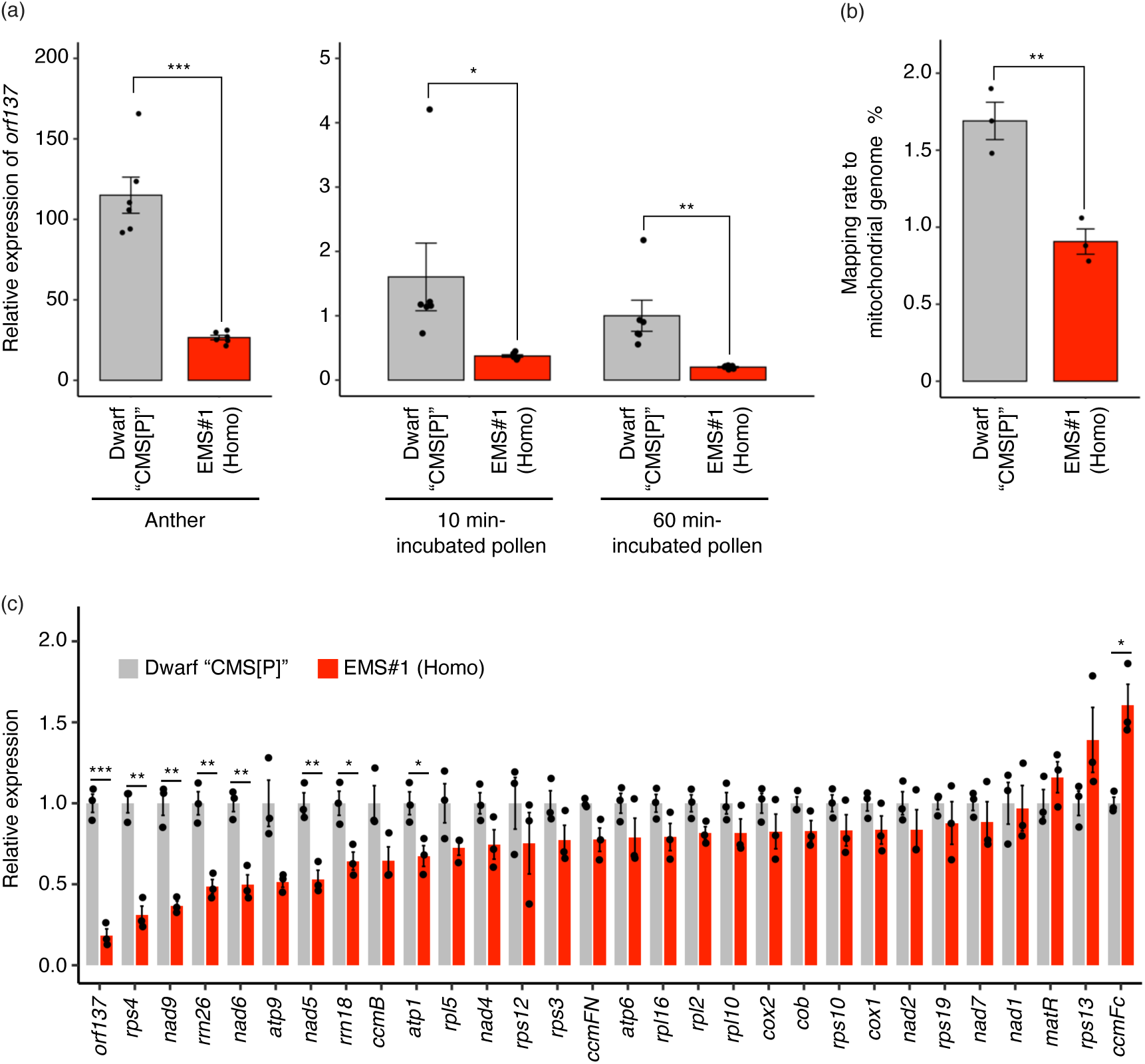
Fertility restoration is associated with a reduced expression level of *orf137*. (a) Relative expression levels of *orf137* measured by RT-qPCR in anthers and pollen incubated for 10 or 60 minutes from Dwarf “CMS[P]” and homozygous EMS#1. Expression levels are normalized to that in 60 min-incubated pollen of Dwarf “CMS[P]” (set to 1). The tomato *actin* gene (*Solyc04g011500*) was used as an internal control. Statistical significance was determined using Student’s t-test. ***P < 0.001; **P < 0.01; *P < 0.05, n = 6. (b) (b) Mapping rate of RNA-Seq reads to the mitochondrial genome in anthers of Dwarf “CMS[P]” and homozygous EMS#1. Statistical significance was determined using Student’s t-test. **P < 0.01, n = 3. (c) Relative expression levels of mitochondrial genes in anthers of Dwarf “CMS[P]” and homozygous EMS#1, as determined by RNA-Seq analysis. The expression levels of each gene in Dwarf “CMS[P]” were set to 1. To minimize the effect of variability, genes with an average count below 50 in Dwarf “CMS[P]” were excluded from the analysis. Statistical significance was determined using Student’s t-test. ***P < 0.001; **P < 0.01; *P < 0.05, n = 3.

### Application of rpoTm mutation in tomato F_1_ hybrid breeding

We investigated whether a heterozygous mutation in EMS#1 could induce sufficient fruit formation in F_1_ plants. To this end, we used three F_1_ plants; F_1_ (P-M) generated by cross-pollination between tomato cultivar “P” and “Micro-Tom”; F_1_ (CMS-M) generated by cross-pollination between “CMS[P]” and “Micro-Tom”; and F_1_ (CMS-*Rf*) generated by cross-pollination between “CMS[P]” and EMS#1. F_1_ (P-M) did not have *orf137* on its mitochondrial genome, whereas F_1_ (CMS-M) and F_1_ (CMS-*Rf*) did. F_1_ (P-M) and F_1_ (CMS-M) did not have *Rf* gene in the nuclear genome, whereas F_1_ (CMS-*Rf*) had a gene derived from EMS#1 as a heterozygote. The percentage of fruits with seeds was 0% in F_1_ (CMS-M) but 100% in F_1_ (P-M) and F_1_ (CMS-*Rf*). In F_1_ (CMS-M), parthenocarpy was observed, and seedless fruits were produced (Figure 6a). Fruits of F_1_ (P-M) and F_1_ (CMS-*Rf*) had similar sizes and weights, which were significantly higher than those of F_1_ (CMS-M). In contrast, the number of seeds in F_1_ fruits tended to decrease in F_1_ (CMS-*Rf*) compared with that in F_1_ (P-M) fruits (Figure 6b). These results demonstrate that the heterozygous mutation in EMS#1 tends to result in fewer seeds in the fruit than the WT F_1_ plants but can induce a 100% fruit-setting rate and sufficient fruit formation.

**Figure 6.**
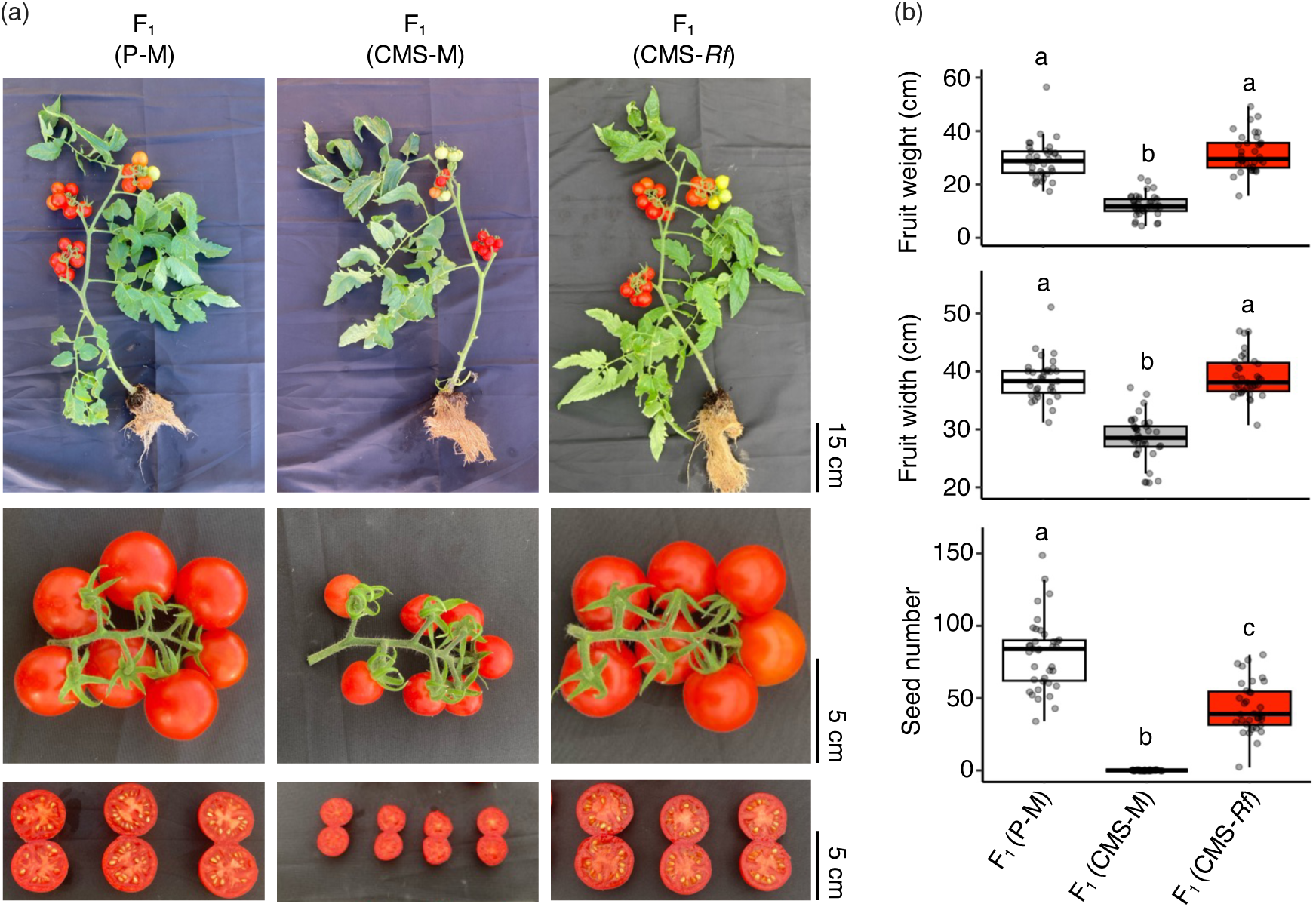
Evaluation of the efficacy of the EMS#1 mutation in F_1_ hybrid breeding. (a) Representative images of three F_1_ hybrid lines: F_1_ (P-M), F_1_ (CMS-M), and F_1_ (CMS-*Rf*). F_1_ (CMS-M) exhibits parthenocarpy and produces seedless fruits. The top row shows whole plants, the middle row displays fruit clusters, and the bottom row presents cross-sections of representative fruits. (b) Comparison of fruit weight, fruit width, and seed number among the three F_1_ hybrid lines. Box plots represent the interquartile range (IQR), black center lines indicate the medians, and whiskers extend to 1.5× IQR. Individual data points represent values from more than 35 independent fruits. Different letters indicate statistically significant differences (P < 0.05, Tukey’s HSD test).

## Discussion

Mutagenic breeding is an important technique for the development of new plant varieties. As the mutations created by EMS are generally heterozygous and recessive in the M_1_ generation, phenotypes are typically investigated in the M_2_ or later generations in mutagenesis breeding (Fonseca *et al*., 2022; Garcia *et al*., 2016). By contrast, we successfully developed novel tomato *Rf* lines at the M_1_ generation by EMS mutagenesis to tomato CMS seeds. There were two reasons for the development of suppressor mutants in the M_1_ generation. (1) The tomato CMS line is a gametophytic CMS in which fertility restoration is determined by the pollen genotype. Although the mutations created by EMS are heterozygous in the M_1_ generation, 50% of the pollen has a mutation that may appear as a phenotype. (2) In this study, we found that the loss of function of *SlRPOTm* was associated with fertility restoration in the tomato CMS line. Most mutations created by EMS mutagenesis are loss-of-function mutations, whereas gain-of-function mutations are extremely rare (Chen *et al*., 2023). Therefore, loss-of-function mutations that restore fertility in CMS are considered compatible with EMS mutagenesis. In addition to the results of the current study, there are other reports of suppressor mutants from CMS seeds. *RFL24* encoding *PPR protein* was identified as an *Rf* gene in the *Arabidopsis* CMS line possessing *orf117Sha*. Furthermore, several suppressor mutants (or called fertile revertants) were isolated by the EMS treatment of *Arabidopsis* CMS lines, and three *rfl24* mutants were obtained by sequencing *RFL24* (Durand *et al*., 2021). In CMS-S maize, suppressor mutants have been isolated from transposon-active lines. Sequencing of the transposon-flanking regions detected transposon insertions in the nuclear genes encoding *RPL6* and *RPL14* in two independent suppressor mutants. It has been confirmed that the loss of function of *RPL6* and *RPL14* restores fertility in CMS-S maize (Gabay-Laughnan *et al*., 2018; Wang *et al*., 2024). *Arabidopsis* CMS lines possessing *orf117Sha* and CMS-S maize are both gametophytic CMS (Bucher, 1961; Gobron *et al*., 2013), similar to the tomato CMS line used in the present study. Therefore, this study demonstrates that mutagenesis breeding can be used to develop novel *Rf* lines for gametophytic CMS, which is expected to be applied to a wide range of plant species in the future. In addition, most *Rf* genes reported to date are derived from the nuclear genomes of wild species and restore fertility as functional genes (Chen and Liu, 2014). In contrast, the *Rf* gene identified in the current study induced fertility restoration through loss of function and was successfully validated using the CRISPR-Cas9 system (Figure 3). In summary, genome editing technology can be used to introduce *rpoTm* mutations into many tomato varieties, facilitating rapid and simple creation of tomato *Rf* lines. As a target mutation of genome editing, an intronic mutation like the one observed in EMS#1 may be preferable, as it confers strong fertility restoration while causing minimal inhibition of embryo development. The CRISPR-Cas9 cytidine deaminase fusion (Shimatani *et al*., 2017), which is capable of C/G-to-T/A substitution, can be used to reproduce the intronic mutation of EMS#1 in other tomato varieties.

In *Arabidopsis*, loss of function of *AtRPOTm* results in embryo lethality owing to the disruption of embryogenesis, and homozygous mutants cannot be acquired. *AtRPOTm* is a major RNA polymerase involved in mitochondrial gene transcription (Kühn *et al*., 2009, 2007; Tan *et al*., 2010). In tomatoes, we were unable to obtain homozygous mutants in EMS#7, 9, and 11 that caused a functional loss of Sl*RPOTm*, such as nonsense and splicing mutations (Table 1, Figure 3a). The results of the RNA-Seq analysis suggested that *SlRPOTm* is involved in the transcription of most mitochondrial genes (Figure 5c), and functional defects in *SlRPOTm* likely cause embryo lethality in tomatoes. In contrast, homozygous mutants with intronic mutations were obtained from EMS #1 (Table 1, Figure 3a). In the mutant with a homozygous intronic mutation, *SlRPOTm* mRNA with eight bases was produced, whereas WT mRNA was also generated (Figure S4). In other words, it is possible to obtain homozygous individuals without complete embryo lethal phenotypes by inducing partial suppression of *RPOTm* expression, rather than the complete loss of function of this gene. The acquisition of the *rpoTm* homozygous mutant allowed further experiments, as homozygous mutants are essential for the investigation of gene function in diploid plant tissues such as leaves and anthers. The homozygous mutant showed a slight reduction in plant growth rate and an increase in days to flowering by several days (Figure S5). In contrast, anther size and pollen number were unchanged in the homozygous mutant, and pollen germination was induced in the cytoplasmic background of CMS (Figure 4). Although the mRNA levels of most mitochondrial genes were reduced in the homozygous *rpoTm* mutant (Figure 5c), the *rpoTm* mutation did not affect anther and pollen development. However, it has been reported that the number of mitochondria per cell increases with anther development and that mitochondrial mRNA levels also increase, suggesting that anther development is an energy-demanding process (Warmke and Lee, 1978; Huang *et al*., 1994; Smart *et al*., 1994). Although our results differ from those of these reports, it has been shown that *rpoTm* mutations do not affect pollen formation or germination in *Arabidopsis rpoTm* heterozygotes (Tan *et al*., 2010). A more detailed analysis of the relationship between anther and pollen development and the mRNA levels of mitochondrial genes is required.

In the present study, we tested whether the heterozygous *rpoTm* mutation induced sufficient fruit formation in F_1_ plants. The number of seeds per fruit in F_1_ (CMS-*Rf*) was lower than that in normal F_1_ (P-M). Intronic mutations derived from EMS#1 partially inhibit embryogenesis, as shown in Table 1, which may explain the lower seed number. In contrast, the fruit size and weight in F_1_ (CMS-*Rf*) were equivalent to those in normal F_1_ (P-M) (Figure 6). Tran *et al* (2023) proposed a model in which full penetration of pollen tubes initiates fruit set by activating the expression of genes regulating cell division and expansion. We confirmed that *rpoTm* mutation induced pollen germination and pollen tube elongation in the cytoplasmic background of CMS (Figure 2 and 3b), which may trigger sufficient fruit enlargement.

In summary, this study provides a method for developing tomato *Rf* lines using EMS mutagenesis and genome editing. Furthermore, we confirmed for the first time that loss of function of *RPOTm* contributes to fertility restoration in CMS lines. Using genome editing, it is possible to replace various tomato varieties with *Rf* lines, which will promote tomato F_1_ breeding in the future.

## Experimental procedures

### Plant growth and EMS treatment

Dwarf “CMS[P]” was developed from tomato CMS line “CMS[P]” by backcrossing with tomato dwarf cultivar “Micro-Tom” (TOMJPF0001) (Scott *et al*., 1989) in our previous study (Kuwabara *et al*., 2022). All plants were cultivated at 23 °C under a 16/8 h light/dark cycle. EMS treatment was performed according to a previously described protocol (Watanabe *et al*., 2007). EMS at 0.5% or 1% (v/v) was applied to seeds of Dwarf “CMS[P]” for 16 h.

### Pollen germination test

A pollen germination test in a liquid germination medium and on the stigma was performed as described by Kuwabara *et al*., 2022. Pollens were observed using a BX53 microscope (Olympus). The ratios of germinated and abnormal pollen were calculated using four biological replicates from independent experiments with more than 300 pollen grains.

### DNA extraction, PCR, DNA marker and Sanger sequencing analysis

Total DNA was extracted from young leaves using a Maxwell 16 instrument and Maxwell 16 Tissue DNA Purification Kit (Promega). PCR was performed with primer sets listed in Table S2 and KOD FX Neo (Toyobo) using the following thermal cycling conditions: initial denaturation at 94 °C for 2 min; 35 cycles of denaturation at 98 °C for 10 s, annealing at 55 °C for 30 s, and extension at 68 °C for 1 min; and final extension at 68 °C for 5 min. PCR products were digested using the restriction enzymes listed in Table S2. For Sanger sequencing (Eurofins Genomics), the PCR product was diluted 10-fold with water and mixed with the primers (Table S2). Data were visualized using SnapGene Viewer v8.0.2 (https://www.snapgene.com).

### Bulk segregant analysis and whole-genome sequencing

Genomic DNA from fertile and sterile individuals was pooled and sequenced on an Illumina NovaSeq 6000 system (Illumina) in PE150 mode. Raw reads were subjected to quality control and adapter trimming using TrimGalore version 0.6.7 (https://github.com/FelixKrueger/TrimGalore), with the settings –length 20 and –quality 30. The clean reads were aligned to the tomato genome reference SL4.0 (Hosmani *et al*., 2019) using BWA-MEM2 version 2.2.1 (Vasimuddin *et al*., 2019). SNPs were identified using elPrep version 5.1.3, with the options --mark-duplicates and remove-duplicates (Herzeel *et al*., 2021). Only SNPs (not indels) were extracted and filtered as follows: QD < 2.0, QUAL < 30.0, SOR > 4.0, FS > 60.0, MQ < 40.0, MQRankSum < −12.5, ReadPosRankSum < −8.0 using GATK version 4.4.0.0 (McKenna *et al*., 2010). SnpEff version 5.2a (Cingolani *et al*., 2012) was used to annotate the SNPs using annotation data for tomato ITAG4.0 (Hosmani *et al*., 2019). Visualization of ΔSNP-index was performed using QTLseqr version 0.7.5.2 (Mansfeld and Grumet, 2018) with default parameters.

### CRISPR-Cas9 construct and tomato transformation

Two guide RNAs for genome editing of *SlRPOTm* genes were determined using CRISPR-P 2.0 (Liu *et al*., 2017), and the sequences were “ACTGCGAGAGGCATTCTCAG” and “CTTGGTTCTGATTTATGCCG.” The pDe-Cas9 vector was used, and this plasmid was modified as previously described (Fauser *et al*., 2014; Mikami *et al*., 2015). The constructed vector was introduced into *Agrobacterium tumefaciens* strain GV3101. The *Agrobacterium*-mediated transformation in Dwarf “CMS[P]” was performed as described previously (Sun *et al*., 2006). The transformed plants were selected using kanamycin.

### Gene expression analysis

Total RNA was extracted from the anthers and incubated with pollen using the RNeasy Plant Mini Kit (Qiagen). The extracted RNA was treated with RNase-free DNase (Qiagen) and used for sequence library preparation using a TruSeq Stranded mRNA Library Prep Kit (Illumina), in which random hexamers were used as primers for reverse transcription. The resultant library was sequenced on a DNBSEQ G400 instrument (MGI Tech) to generate 100-bp paired-end reads. Raw reads were subjected to quality control and adapter trimming using TrimGalore version 0.6.7 (https://github.com/FelixKrueger/TrimGalore), with settings of –length 20 and –quality 30. Clean reads were mapped to the mitochondrial genome reference of “CMS[P]” (accession number LC613119-LC613123 in DDBJ database) (Kuwabara *et al*., 2021) using HISAT2 version 2.2.1 (Kim *et al*., 2015). Mitochondrial gene annotation of the “CMS[P]” genome was performed using the online tool GeSeq (Tillich *et al*., 2017) based on the tomato and potato reference genomes (GenBank accession numbers: MF034192, MF034193, MN104801-MN104803, and MN114537-MN114539). To avoid cross-mapped plastid reads, regions identical to plastid DNA of “CMS[P]” (accession number LC613092-LC613094 in the DDBJ database) were detected using BLAST (Ye *et al*., 2006) with parameters of -perc_identity 95 -evalue 1e-5 and masked using N (A or T or G or C). The read counts mapped to the masked mitochondrial genes were calculated using RSEM (Li and Dewey, 2011). The relative expression levels of each gene were calculated by dividing the mapped read counts by the total number of sequence reads for each sample.

For RT-qPCR, cDNA was synthesized from 800 ng of total RNA using the ReverTra Ace qPCR RT Kit (Toyobo) with random hexamers as primers. RT-qPCR for *orf137* was performed using TB Green Premix Ex Taq II (Tli RNaseH Plus) (Takara Bio) and CFX96 Real-Time PCR Detection System (Bio-Rad) according to previously described procedures (Shinozaki *et al*., 2015) with six biological and two technical repeats. The delta-delta CT method (Pfaffl, 2001) was used to calculate gene expression levels. The tomato a*ctin* gene (accession no. *Solyc04g011500*) was used as an internal control (Zhang *et al*., 2008). The specific primers used for each gene are listed in Table S2.

### Subcellular localization of SlRPOTm

The full-length cDNA fragment of *SlRPOTm* was amplified from the genomic DNA of *S. lycopersicum* ‘Micro-Tom’. The amplified fragment was inserted into the pENTR™ /D-TOPO™ Vector (Thermo Fisher Scientific) by In-Fusion HD Cloning Kit (Takara Bio). The super folder green fluorescent protein (sfGFP) fusion vector was constructed as follows: First, the pUGW_35SsGFPHSP vector was generated by inserting a PCR-amplified Gateway cassette from pDEST35S_sfG-GW_HSPT (Ezura *et al*., 2022) into the SmaI-digested p35S_GFP_HSPG vector (Hu *et al*., 2022). Then, the sfGFP sequence was amplified from a pDEST35S_sfG-GW_HSPT (Ezura *et al*., 2022), while the vector backbone sequence with 35S promoter, Gateway cassette, and HSP terminator was amplified from pUGW_35SsGFPHSP. These amplified fragments were subsequently assembled using NEBuilder HiFi DNA Assembly (New England BioLabs) to generate a transient expression vector, pUGW_sfGFP_HSPT. The subcloned *SlRPOTm* sequence was transferred into the vector by Gateway LR reaction (Thermo Fisher Scientific). Primers used in plasmid construction are listed in Table S2.

Protoplasts were isolated from leaves of *N. benthamiana* and transfected following a previously described protocol (Lin *et al*., 2018) with some minor modifications as follows: To prepare protoplasts, leaves were sliced and incubated in a 25-mL tube containing 10 mL digestion solution (1.5 % (w/v) Cellulase “Onozuka” R-10 (Yakult), 0.75 % (w/v) macerozyme R10 (Yaklut), 10 mM 2-mercaptoethanol and 0.6 M mannitol, pH5.7) for two hours with gentle swinging. After incubation, protoplasts were released by agitating the tube at 50 rpm for 5 minutes. The digested solution was filtered through 70-µm cell strainers into a 50-mL tube. Then, the 25-mL tube was rinsed with 10 mL pre-cooled W5 solution (150 mM NaCl, 125 mM CaCl_2_, 5 mM KCl, 2 mM MES, and 5 mM glucose), and the rinse was filtered again with the cell strainer into the same 50-mL tube. Protoplasts were collected by centrifugation at 200 × g and washed twice with 20 mL W5 solution. The protoplast pellet was resuspended in 10 mL of W5 solution and incubated on ice for 15 minutes. Meanwhile, plasmid DNA was prepared in round-bottom 2.0-mL tubes. After incubation, protoplasts were collected again by centrifugation at 200 x g for 3 minutes, and the supernatant was removed. The pellet was then resuspended in 2.0 mL pre-cooled MMG solution (0.6 M mannitol, 15 mM MgCl_2_, 4 mM MES, pH5.7). Protoplasts were counted using a hemocytometer under a microscope, and cell density was adjusted to 2.5-5.0 × 10^5^ cells/mL with pre-cooled MMG solution. For transfection, protoplasts were subjected to PEG-mediated transfection. A 200-µL aliquot of protoplast suspension was added to each 2.0-mL tube containing at least 4 µg of plasmid DNA in 20 µL and gently mixed by inverting 5-10 times. An equal volume of PEG solution (40% (w/v) PEG 4000, 0.6 M mannitol, 0.1 M CaCl_2_) was then added and gently mixed by inverting 5 −10 times. The mixture was incubated at room temperature in the dark for 15 minutes. To terminate the reaction, 1 mL of W5 solution was added and mixed well. The transfected protoplasts were collected by centrifugation at 200 x g for 3 minutes, washed twice, and resuspended in fresh W5 solution. Finally, protoplasts were incubated overnight at room temperature in the dark before observation.

For mitochondria staining, protoplasts were treated with 200 nM MitoTracker Red CMXRos (Invitrogen) for 30 minutes, washed three times with W5 solution, and then observed using a confocal laser scanning microscope (LSM700, Zeiss). Fluorescence was detected as follows: GFP was excited at 488 nm, with detection between 490 and 555 nm; MitoTracker Red was excited at 555 nm, with detection between 505 and 600 nm; and chlorophyll autofluorescence was excited at 639 nm, with detection between 657 and 726 nm.

### CR-RT-PCR to determine 5’ and 3’ UTR region of orf137

CR-RT-PCR was performed as described previously (Forner *et al*., 2007). Briefly, 4 µg total RNA extracted from anthers of “CMS[P]” was circularized with T4 RNA Ligase (Takara Bio). Reverse transcription of the circularized RNA was conducted using the primers orf137_CR-RT (Table S2) and SuperScript III Reverse Transcriptase (Invitrogen). Nested PCR was performed using primer sets of orf137_CR-RT_Fw1 and orf137_CR-RT_Rv1 for the first PCR, and orf137_CR-RT_Fw2 and orf137_CR-RT_Rv2 for the second PCR and Ex-Taq polymerase (Takara Bio) using the following thermal cycling conditions: initial denaturation at 94 °C for 1 min; 35 cycles of denaturation at 98 °C for 10 s, annealing at 55 °C for 30 s, and extension at 72 °C for 1 min; and final extension at 72 °C for 5 min. After electrophoresis on a 1% (w/v) agarose gel, the PCR products were purified using a FastGene Gel/PCR Extraction Kit (Nippon Genetics, Tokyo, Japan). After cloning the PCR products into the pGEM-T easy vector (Promega), the sequences of 28 clones were determined using Sanger sequencing (Eurofins Genomics).

## Data availability

Raw read data for BSA was deposited in the Sequence Read Archive (SRA) database of the DNA Data Bank of Japan (DDBJ) under the BioProject accession number PRJDB20274. Raw read data for RNA-Seq was deposited under the BioProject accession number PRJDB20280.

## Author contributions

T.A., K.S., K.T., and K.K. conceived and coordinated the study. K.K., R.N., B.V., and K.E. performed most of the experiments. K.S. and K.K. performed sequencing analysis. K.S., K.T., K.E., and K.K. wrote the manuscript.

## Acknowledgments

We thank all the technical and administrative members of the T-PIRC Center at the University of Tsukuba and Kazusa DNA Research Institute. We are also grateful to Masaki Endo for kindly providing the CRISPR-Cas9 plasmids and to Dr. Shingo Sakamoto (National Institute of Advanced Industrial Science and Technology, Japan) for kindly providing the pUGW_35SsGFPHSP vector. Computations were partially performed on the NIG supercomputer at ROIS National Institute of Genetics. Micro-Tom (TOMJPF0001) was provided from National BioResource Project Tomato (NBRP tomato). We would like to thank Editage (www.editage.jp) for English language editing. This research was supported by the Project of the NARO Bio-oriented Technology Research Advancement Institution (Research Program on Development of Innovative Technology, Grant Number: JPJ007097) to K.S. and T. A. and JSPS Research Fellowships for Young Researchers (Grant Number: 21J20479) to K.K, and the Kazusa DNA Research Institute Foundation to K.S.

## Conflict of interest disclosure

The authors have no competing interests to declare.

## Short legends for Supporting Information

**Table S1** Summary of RNA-Seq analysis.

**Table S2** Primers used in this study.

**Figure S1** Bulked Segregant Analysis for the fertile and sterile population in EMS#1.

**Figure S2** Subcellular localization of SlRPOTm in *Nicotiana benthamiana* protoplasts.

**Figure S3** Evidence for homozygous mutation in EMS#1.

**Figure S4** Evaluation of the intronic mutation in EMS#1 and its effect on *SlRPOTm* splicing.

**Figure S5** Effects of heterozygous and homozygous EMS#1 mutations on vegetative tissues.

**Figure S6** Determination of the 5’ and 3’ UTR regions of *orf137* by CR-RT-PCR.

## Notes

### Competing Interest Statement

The authors have declared no competing interest.

