## supplemental Files for "Artificial mutations in the nuclear gene encoding mitochondrial RNA polymerase restore pollen fertility in cytoplasmic male sterile tomato"

**Table S1** Summary of RNA-Seq analysis.

| Line | Tissue | Repeat | Total pairs | Aligned concordantly 1 time | Aligned concordantly >1 times | Overall alignment rate % |
|---|---|---|---|---|---|---|
| Dwarf “CMS[P]” | Anther | 1 | 23653473 | 273616 | 125278 | 1.69 |
| Dwarf “CMS[P]” | Anther | 2 | 25135832 | 327827 | 150239 | 1.90 |
| Dwarf “CMS[P]” | Anther | 3 | 21204417 | 215849 | 98079 | 1.48 |
| EMS#1 (homo) | Anther | 1 | 21977271 | 145698 | 88268 | 1.06 |
| EMS#1 (homo) | Anther | 2 | 23193966 | 112544 | 69141 | 0.78 |
| EMS#1 (homo) | Anther | 3 | 23955898 | 131251 | 80301 | 0.88 |
| Dwarf “CMS[P]” | 10 min-incubated pollen | 1 | 23214276 | 3613 | 1812 | 0.02 |
| Dwarf “CMS[P]” | 10 min-incubated pollen | 2 | 22717653 | 11083 | 5940 | 0.07 |
| Dwarf “CMS[P]” | 10 min-incubated pollen | 3 | 24737850 | 3648 | 1936 | 0.02 |
| EMS#1 (homo) | 10 min-incubated pollen | 1 | 22676570 | 2895 | 1849 | 0.02 |
| EMS#1 (homo) | 10 min-incubated pollen | 2 | 19158037 | 2012 | 1412 | 0.02 |
| EMS#1 (homo) | 10 min-incubated pollen | 3 | 18896669 | 1918 | 1313 | 0.02 |
| Dwarf “CMS[P]” | 60 min-incubated pollen | 1 | 23727617 | 3764 | 1929 | 0.02 |
| Dwarf “CMS[P]” | 60 min-incubated pollen | 2 | 24443592 | 9078 | 5495 | 0.06 |
| Dwarf “CMS[P]” | 60 min-incubated pollen | 3 | 22677189 | 3734 | 1955 | 0.03 |
| EMS#1 (homo) | 60 min-incubated pollen | 1 | 25013581 | 2345 | 1593 | 0.02 |
| EMS#1 (homo) | 60 min-incubated pollen | 2 | 18499882 | 1566 | 994 | 0.01 |
| EMS#1 (homo) | 60 min-incubated pollen | 3 | 19679214 | 1401 | 954 | 0.01 |

**Table S2** Primers used in this study.

| Name | Sequence (5' to 3') | Application |
|---|---|---|
| EMS#1_CAPS_Fw | TCCACTAGCCCATATTCTGTCTAAT | CAPS marker for EMS#1 (PCR product is digested by HaeⅢ) |
| EMS#1_CAPS_Rv | CTCAGAACCATGTATGGATTGGTT | CAPS marker for EMS#1 (PCR product is digested by HaeIII) |
| EMS#7_CAPS_Fw | GTGAATGGGCATATGAGATATAGAGGT | CAPS marker for EMS#7 (PCR product is digested by HphⅠ) |
| EMS#7_CAPS_Rv | CTGATTTATGCCGTGGCATCC | CAPS marker for EMS#7 (PCR product is digested by HphⅠ) |
| EMS#9_CAPS_Fw | GAATTTGATCAAACTTCCTGCAGC | CAPS marker for EMS#9 (PCR product is digested by HphⅠ) |
| EMS#9_CAPS_Rv | CCATTGAATAAGCTGTCTTTGTTTGA | CAPS marker for EMS#9 (PCR product is digested by HphⅠ) |
| EMS#11_dCAPS_Fw | ACACCACCACCATATACATTTGCTAGATGTATCTTCAGCCAGCGTAAtC† | CAPS marker for EMS#11 (PCR product is digested by XbaⅠ) |
| EMS#11_dCAPS_Rv | ACTGCGAGAGGCATTCTCAG | CAPS marker for EMS#11 (PCR product is digested by XbaⅠ) |
| EMS#1_exon6_Fw | GTATGACAGAGGTGCATACTTATTTTTACCA | RT-PCR and Sanger-sequencing for an intronic mutaion in EMS#1 |
| EMS#1_exon7_Rv | ATCTTCCCTGTCGACTAAATCAGCAA | RT-PCR and Sanger-sequencing for an intronic mutaion in EMS#1 |
| NPTⅡ_Fw | ATGATTGAACAAGATGGATTGCAC | PCR for *NPTⅡ* |
| NPTⅡ_Rv | TCAGAAGAACTCGTCAAGAAGGCG | PCR for *NPTⅡ* |
| gRNA_Target1_Fw | ATTGACTGCGAGAGGCATTCTCAG | Construction of CRISPR-Cas9 vector |
| gRNA_Target1_Rv | AAACCTGAGAATGCCTCTCGCAGT | Construction of CRISPR-Cas9 vector |
| gRNA_Target2_Fw | ATTGCTTGGTTCTGATTTATGCCG | Construction of CRISPR-Cas9 vector |
| gRNA_Target2_Rv | AAACCGGCATAAATCAGAACCAAG | Construction of CRISPR-Cas9 vector |
| CR_sequence_Fw | GCCGTCTTGCTGATTTAGTCGA | Sequence confirmation of CRISPR-Cas9-mediated mutants |
| CR_sequence_Rv | ATATCCTCCACATGGTTTTCACTG | Sequence confirmation of CRISPR-Cas9-mediated mutants |
| orf137_RT-qPCR_Fw | GGTACATCGCTTTCTCTGTGTC | RT-qPCR for *orf137* |
| orf137_RT-qPCR_Rv | GAGTTTCGTCCCCGTCTTAATTTTC | RT-qPCR for *orf137* |
| Actin_RTqPCR_Fw | GCGAGAAATTGTCAGGGACGT | RT-qPCR for *Actin* |
| Actin_RTqPCR_Rv | TGCCCATCTGGGAGCTCAT | RT-qPCR for *Actin* |
| SlRPOTm_InFusion_Fw | GCAGGCTCCGCGGCCGCCACCATGTGGAGATACATATCAAAACAAGTTT | Amplification of *SlRPOTm* coding region for InFusion cloning |
| SlRPOTm_InFusion_Rv | AGCTGGGTCGGCGCGCCCGTTGAAAAAATAGGGAGATTCAAGAACT | Amplification of *SlRPOTm* coding region for InFusion cloning |
| pENTR/D-TOPO_InFusion_Fw | CGCGCCGACCCAGCTTTCTTGTACAAAGTT | Inverse PCR of pENTR™ /D-TOPO™ vector for InFusion cloning |
| pENTR/D-TOPO_InFusion_Rv | GGCCGCGGAGCCTGCTTTTTTGTACAAAGT | Inverse PCR of pENTR™ /D-TOPO™ vector for InFusion cloning |
| sfGFP_P1 | ATGGTGAGCAAGGGCGAGGAG | Construction of pUGW_sfGFP_HSPT vector |
| sfGFP_P2 | TTTATACAGCTCGTCCATGCCGAGAGTGATCCC | Construction of pUGW_sfGFP_HSPT vector |
| vector_P1 | CTCCTCGCCCTTGCTCACCATCCCCATCACCACTTTGTACAAGAAAGCT | Construction of pUGW_sfGFP_HSPT vector |
| vector_P2 | GGGATCACTCTCGGCATGGACGAGCTGTACAAGTAAGTCGACGAGCTCATATGAAGATGAAGA | Construction of pUGW_sfGFP_HSPT vector |
| GW_cassette_P1 | GATCACAAGTTTGTACAAAAAAGC | Construction of pUGW_sfGFP_HSPT vector |
| GW_cassette_P2 | CATCACCACTTTGTACAAGAAAGCTG | Construction of pUGW_sfGFP_HSPT vector |
| orf137_CR-RT | TAGGCGATCTCAGGTTGACCG | cDNA synthesis of circularized RNA |
| orf137_CR-RT_Fw1 | AGGATTAACGCAAATGTATATTGATC | 1st PCR in the cR-RT-PCR experiment |
| orf137_CR-RT_Rv1 | GGTCGAGTTAAGGTGAAGACTG | 1st PCR in the cR-RT-PCR experiment |
| orf137_CR-RT_Fw2 | GGAGTAACGAGTAAGTACTTTTCTT | 2nd PCR in the cR-RT-PCR experiment |
| orf137_CR-RT_Rv2 | ATCACGGAACACTTTCTATATCTAAG | 2nd PCR in the cR-RT-PCR experiment |

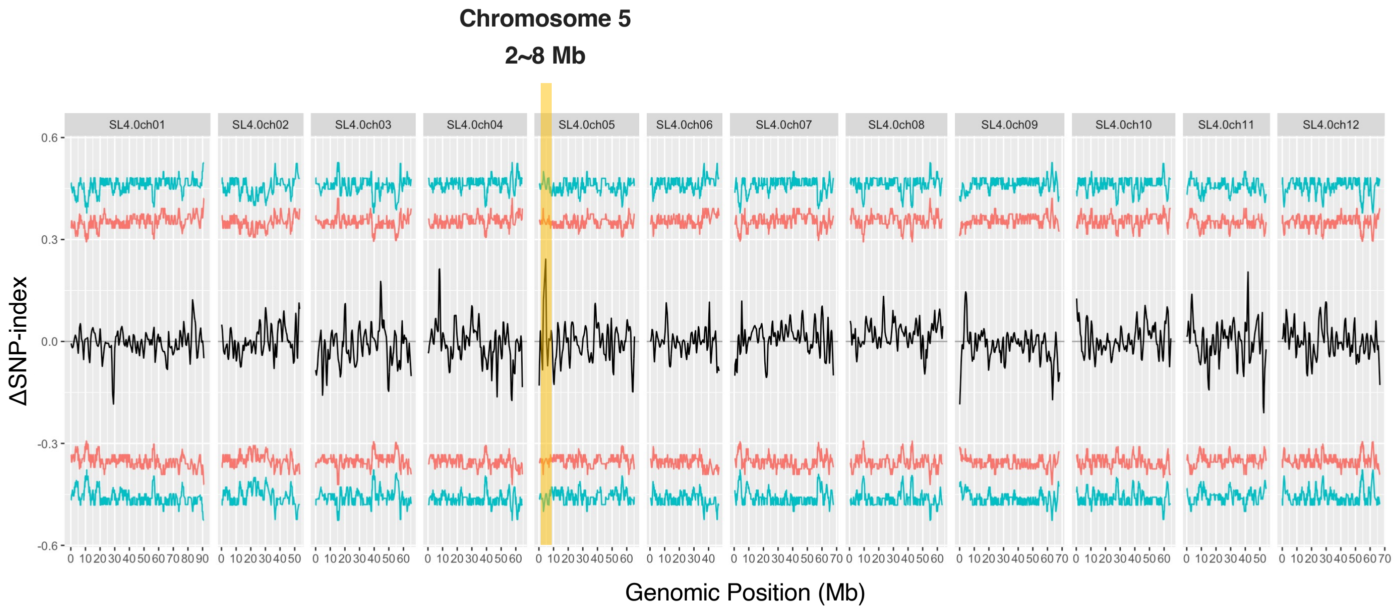


**Figure S1** Bulked Segregant Analysis for the fertile and sterile population in EMS#1. Vertical indicates the ΔSNP-index between fertile and sterile bulk calculated in a 1 Mb sliding window. Horizontal indicates the position of chromosomes in tomato reference genome, Heinz 1706 (SL4.0). Red and blue lines indicate 95% and 99% confidence intervals, respectively. The orange shaded area is region (SL4.0ch05: 2~8 Mb) that showed relatively high ΔSNP-index and was used for further analysis.

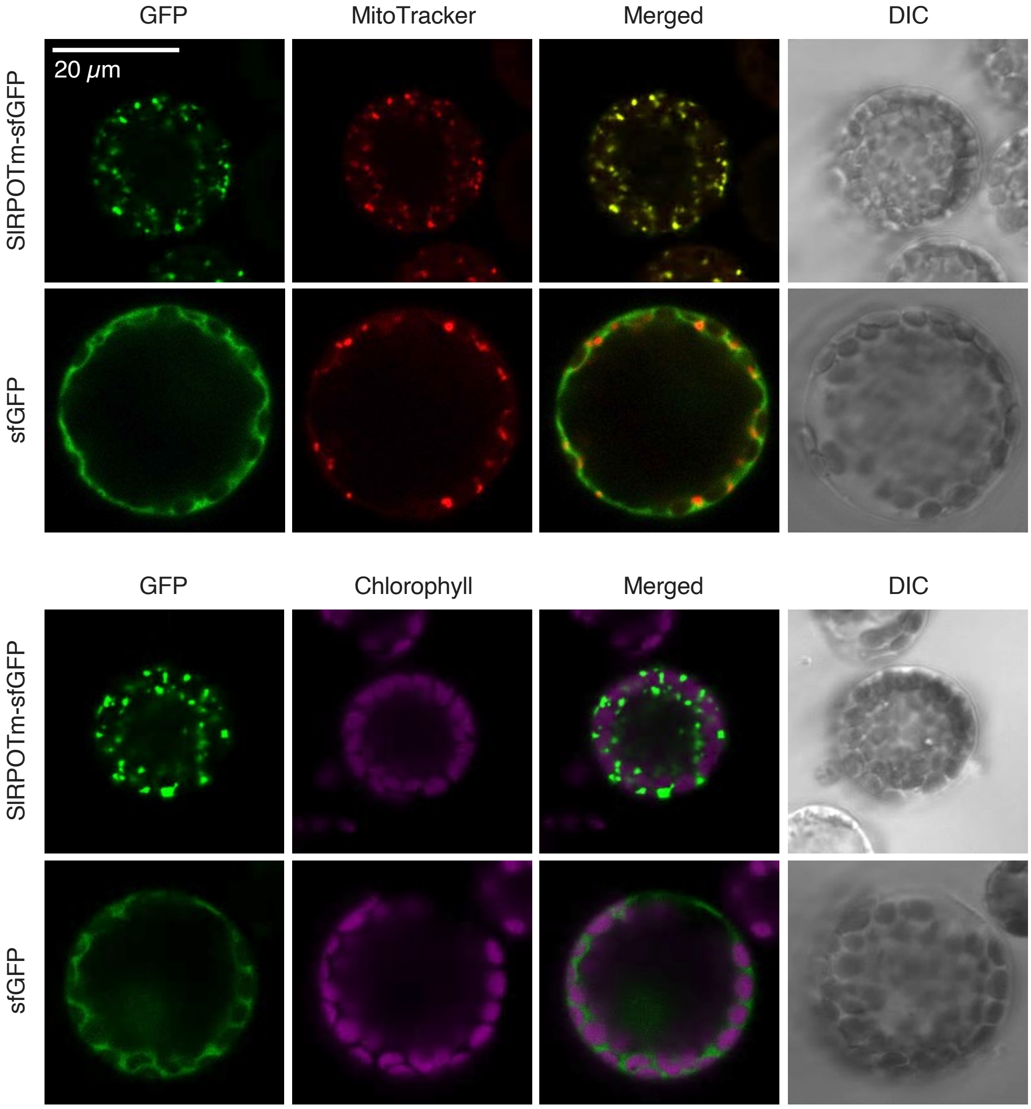


**Figure S2** Subcellular localization of SlRPOTm in *Nicotiana benthamiana* protoplasts. sfGFP was fused to the C-terminus of SlRPOTm and transiently expressed in *N. benthamiana* protoplasts to investigate its subcellular localization. Fluorescence images show GFP signals (green), MitoTracker signals (red, top panel), and chlorophyll autofluorescence (magenta, bottom panel). Merged images indicate colocalization, and differential interference contrast (DIC) images are provided for structural reference.

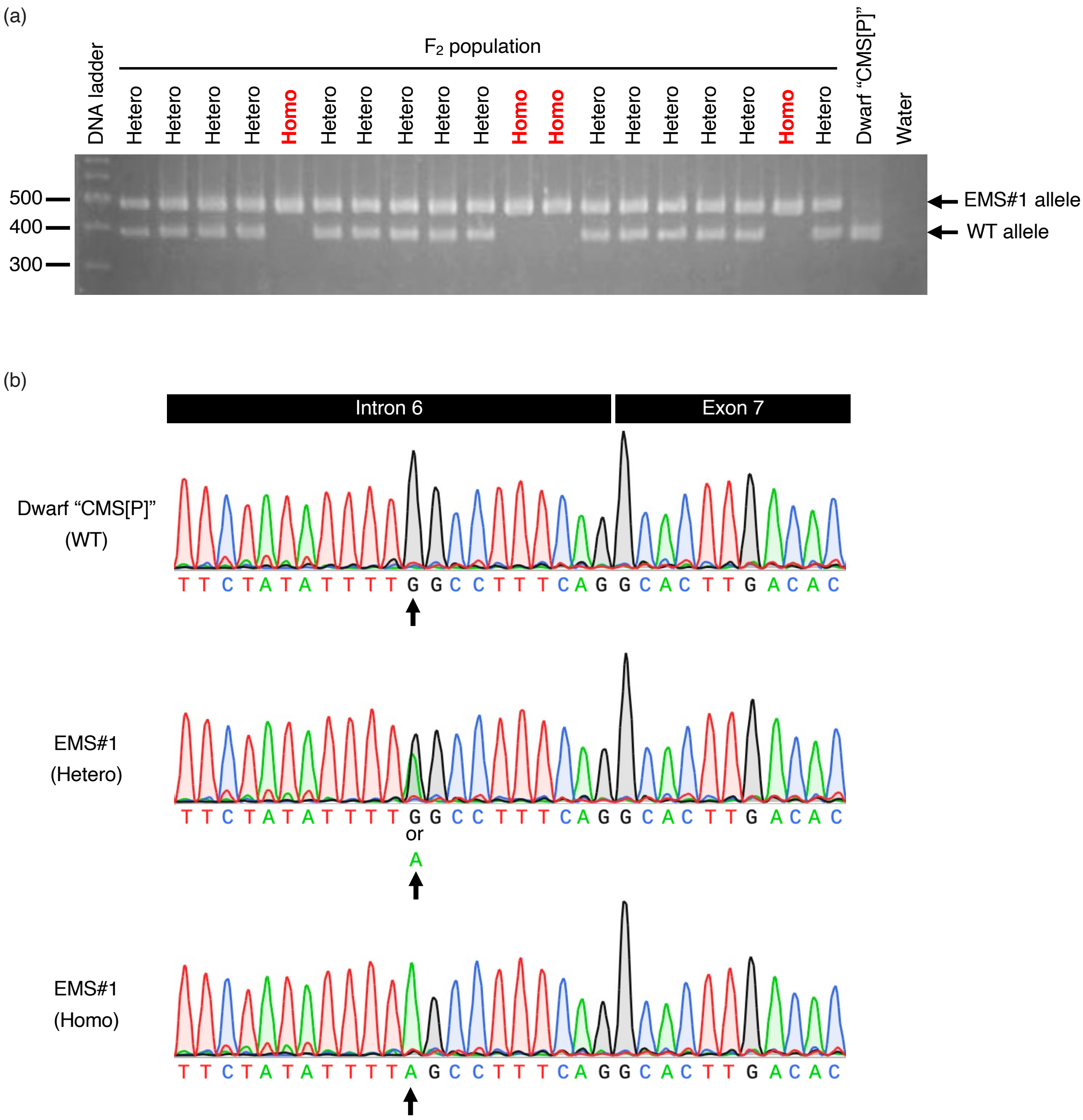


**Figure S3** Evidence for homozygous mutation in EMS#1. (a) Cleaved Amplified Polymorphic Sequence (CAPS) marker analysis of the $BC_2F_2$ population of EMS#1. DNA fragments amplified from individual plants were digested with a restriction enzyme and separated by gel electrophoresis. The presence of both the EMS#1 allele and the wild-type (WT) allele indicates heterozygosity, while the presence of only the EMS#1 allele indicates homozygosity. (b) Sanger sequencing chromatograms showing the mutation site in intron 6 of *SlRPOTm* gene in Dwarf "CMS[P]" (WT), heterozygous EMS#1, and homozygous EMS#1 plants. The black arrow indicates the mutation position (G to A transition in EMS#1).

(a)

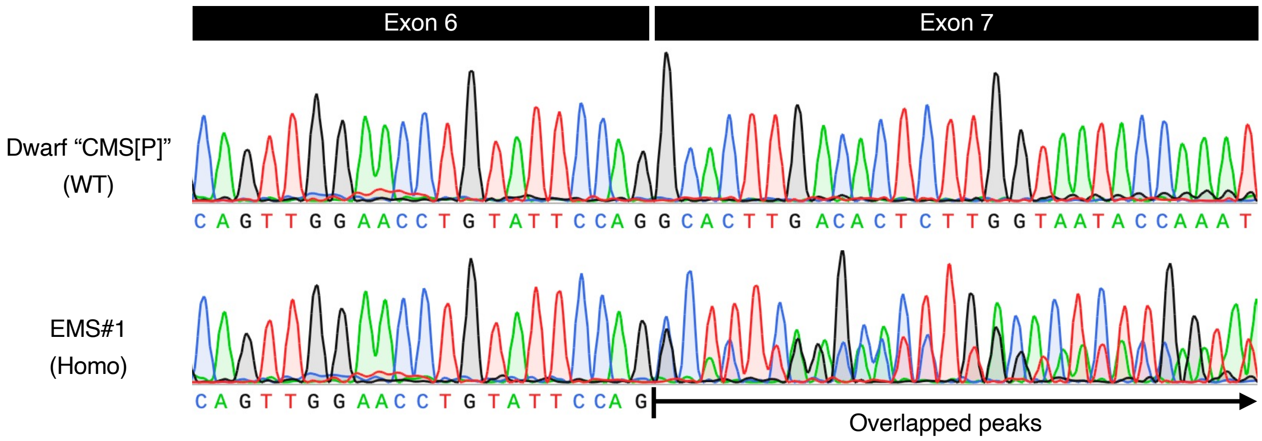


(b)

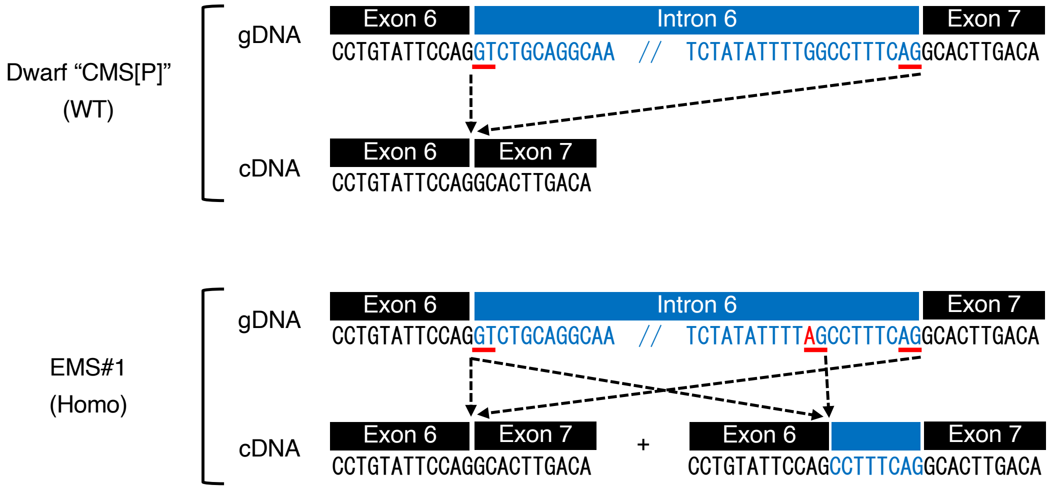


**Figure S4** Evaluation of the intronic mutation in EMS#1 and its effect on *SlRPOTm* splicing. (a) Sanger sequencing chromatograms of RT-PCR products from Dwarf "CMS[P]" (WT) and homozygous EMS#1. Overlapping peaks (black arrow) were detected between exons 6 and 7 in EMS#1, suggesting alternative splicing at this region. (b) Schematic representation of the genomic (gDNA) and cDNA sequences of *SlRPOTm* in Dwarf "CMS[P]" (WT) and homozygous EMS#1. In WT, normal splicing removes intron 6, joining exon 6 to exon 7. In homozygous EMS#1, an intronic G-to-A mutation 10 bp upstream of exon 7 creates a new splicing site "AG." As a result, alternative splicing occurs, generating two transcript variants: one with the original splicing site and another with the newly formed site, leading to an 8 bases insertion in the mature mRNA.

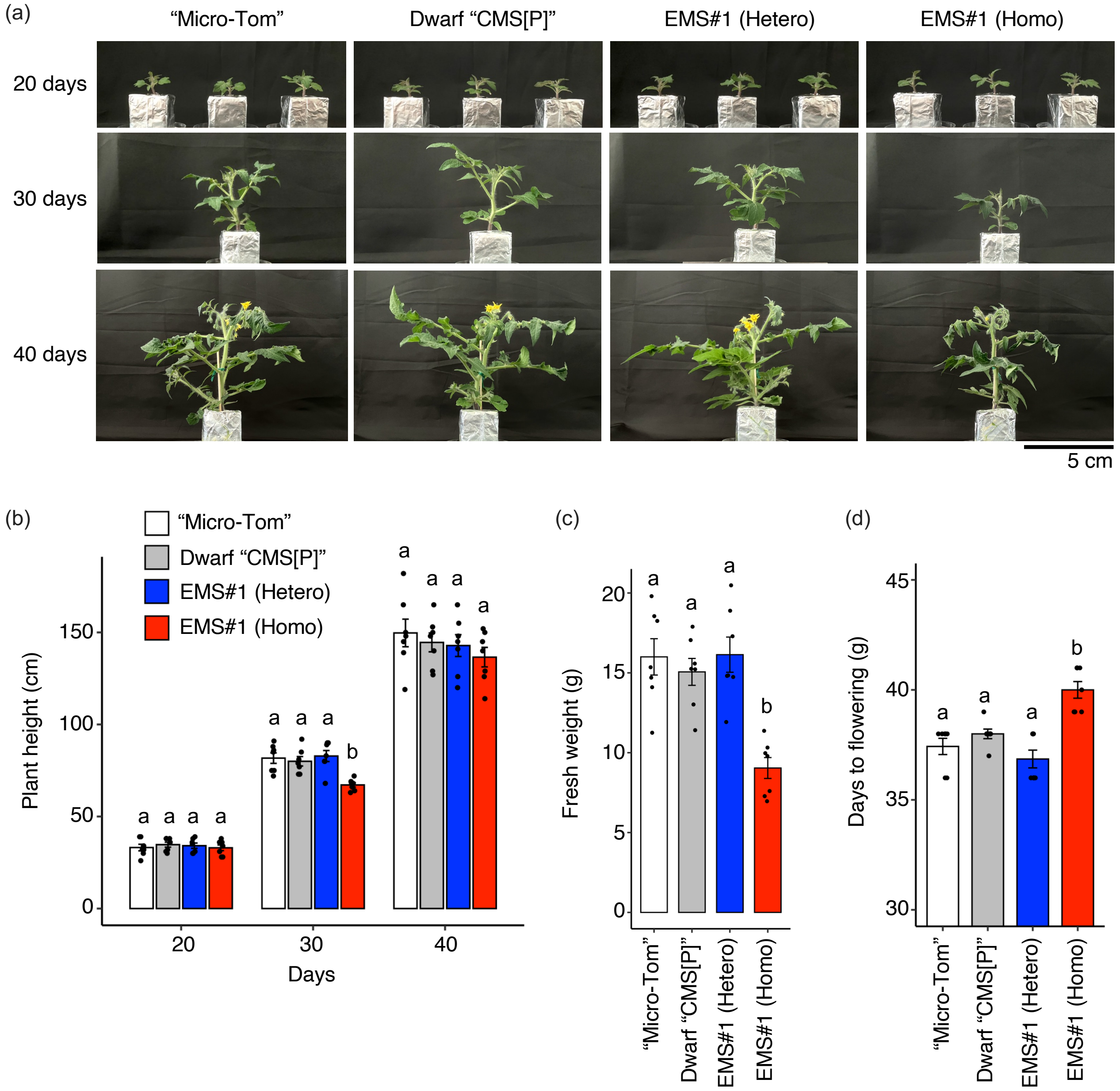


**Figure S5** Effects of heterozygous and homozygous EMS#1 mutations on vegetative tissues. (a) Representative images of plant growth at 20, 30, and 40 days after sowing for WT "Micro-Tom," Dwarf "CMS[P]," heterozygous EMS#1, and homozygous EMS#1 mutants. (b) Plant height measured at 20, 30, and 40 days after sowing (n = 7). (c) Fresh weight of plants at 43 days after sowing (n = 7). (d) Days to flowering for each genotype (n = 7). Different letters indicate statistically significant differences ($P < 0.05$).

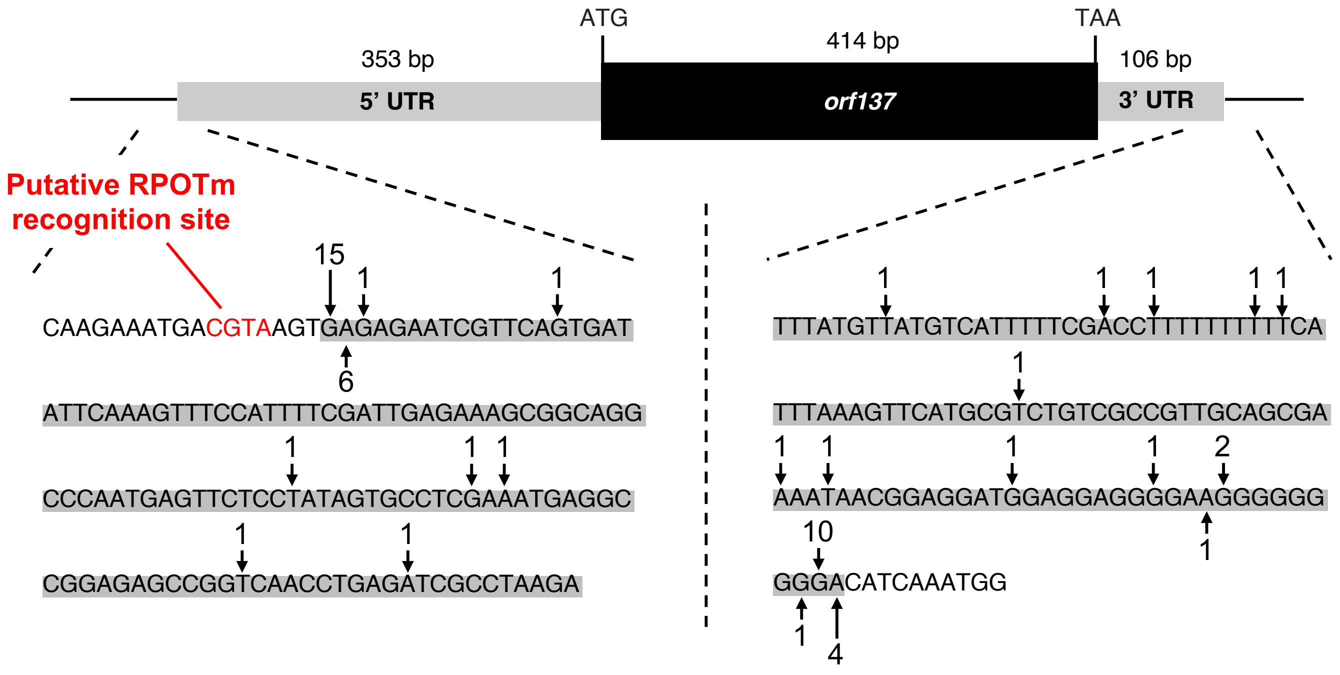


**Figure S6** Determination of the 5’ and 3’ UTR regions of *orf137* by CR-RT-PCR. The numbers shown with black arrows indicate the number of clones obtained by CR-RT-PCR. The nucleotide sequences highlighted with a gray background represent untranslated region (UTR) regions. The 5’ UTR of *orf137* was determined to be 353 bp, while the 3’ UTR was 106 bp by CR-RT-PCR. A putative RPOTm recognition site (CGTA), which is part of the YRTA motif, was identified 4 bp upstream of the 5’ UTR, suggesting that SlRPOTm recognizes this sequence and initiates transcription of *orf137*.
